# Diversity of egg parasitoids of stink bugs in France, with emphasis on Scelionidae parasitizing main Pentatomidae pests

**DOI:** 10.64898/2026.09.25.754315

**Authors:** Alexandre Bout, Sylvie Warot, Lily Cesari, Rachid Hamidi, Claire Caravel, Elijah Talamas, Francesco Tortorici, Nicolas Ris

## Abstract

Stink bugs (Hemiptera Pentatomidae) are major agricultural pests worldwide, including both native and invasive species. In France, for instance, *Halyomorpha halys* has rapidly expanded its range, affecting key crops such as hazelnuts, apples, and vegetables. Meanwhile, several native, polyphagous species — *Nezara viridula*, *Palomena prasina* and, to a lesser extent, *Graphosoma italicum* — are re-emerging and impacting diverse crops because of changes in agricultural practices and climatic conditions.

Egg parasitoids represent promising biological control agents for these pests, be it through conservation, augmentation or introduction biological control. However, their actual potential is not precisely documented due to a lack of ambitious surveys and possible taxonomic issues. This study aims to address this gap by compiling results obtained from various of egg parasitism surveys of Pentatomidae in France and implementing a large-scale DNA Barcoding approach (2000 individuals) for the emerged parasitoids.

Our results indicate a high diversity of egg parasitoids, and detected some taxonomic issues and molecular inconsistencies that warrant further study. Among the three families of egg parasitoids recovered (Encyrtidae, Eupelmide, and Scelionidae), Scelionidae was the most abundant and diverse with nine species of *Trissolcus* (including the two adventive species, *Tr. japonicus* and *Tr. mitsukurii*) and four *Telenomus* species (including two unidentified to species). Scelionidae dominated the parasitoid assemblages associated with most of the Pentatomidae host species, except for *H. halys* for which the eupelmid *Anastatus bifasciatus* was the most common species.

**Key Message:**

- Egg parasitoids assemblages associated with stink bug pests in France remain incompletely documented at the national scale.
- We combined field surveys with large-scale COI barcoding of ∼2,000 parasitoids.
- Scelionidae dominate: 9 *Trissolcus* and 4 *Telenoms* species on most Pentatomidae.
- *Anastatus bifasciatus* (Eupelmidae) prevails on invasive *Halyomorpha halys*.
- Host-specific communities should guide egg-parasitoid choice for biocontrol.

## Introduction

Pentatomid bugs include several highly polyphagous species whose feeding damage affects fruit, nut, vegetable and seed crops, and whose management has become increasingly difficult in both invaded and long-established agroecosystems (Bosco et al. 2018; Leskey & Nielsen, 2018; Mc Pherson, 2018; Conti et al. 2021; Zapponi et al. 2021).

Their spread and pest status are influenced by international trade, accidental introductions, climate warming and changes in crop-protection practices. The brown marmorated stink bug, *Halyomorpha halys* (Stål, 1855), illustrates this trend. Native to East Asia, it has become a major invasive agricultural pest in North America (USA and Canada) in the 1990s (Leskey et al. 2012; Leskey & Nielsen, 2018). In the USA, *H. halys* caused over 37 million dollars in damages to apple orchards in the Mid-Atlantic region in 2010 (Leskey et al. 2012). *H. halys* was detected in Europe during the early 2000s and subsequently reported from several countries including France (Switzerland in 2007; Italy, Germany and France in 2012) (Haye et al. 2014; Cianferoni et al. 2018; Zapponi et al. 2021). In France, risk assessments have highlighted the potential impact of *H. halys* on several fruit, nut and vegetable crops (Haye et al. 2014;). In France, damage to hazelnut orchards increased from 5% in 2017 to 10% in 2023, with local infestation levels exceeding 40% (Hamidi, pers. Comm.).

In addition to recently invasive species, long-established pentatomids may also become more problematic under changing crop-management practices, including the reduced availability or use of broad-spectrum insecticides, and possibly due to climate warming. For example, *Nezara viridula* (Linnaeus, 1758) is a highly polyphagous stink bug, now distributed worldwide in tropical, subtropical, and warm temperate regions. In Europe, including France, its range has expanded over recent decades, and the species continues to extend its distribution northward, likely favoured by climate warming (Kiritani 2006; 2011). In France, *N. viridula* is mainly considered a pest of protected vegetable crops including eggplant (Gard et al. 2022) and tomato (Blancard 2026), but it also occurs in open field crops such as soybean (Ballanger and Jouffret, 1997). *Palomena prasina* (Linnaeus, 1761) remained a key pest of hazelnut orchards until displaced by *H. halys* (Driss et al. 2024; Gomes et al. 2025). *Rhaphigaster nebulosa* Poda, and *Gonocerus acuteangulatus* continue to expand northward and are increasingly reported as secondary pests, where they may cause damage to apples, pears, and other fruit crops (Alkarrat et al. 2020; Beliën et al. 2015; Powell 2020; Hamidi et al. 2022). Finally, other stink bugs, such as *Eurydema* Laporte de Castelnau (1832) and *Graphosoma* Laporte de Castelnau (1833), may be locally relevant in vegetable or seed-production systems, although their economic importance is more context dependent. They also remain useful hosts for documenting parasitoid diversity and host associations within Pentatomidae.

In Europe, and particularly in France, the use of chemical insecticides has become increasingly restricted due to stricter regulatory requirements aimed at reducing their environmental impact and risks to human health. The progressive withdrawal of several active substances and the implementation of integrated pest management (IPM) strategies have encouraged the development of more sustainable crop protection approaches.

Consequently, research efforts are increasingly focused on alternative control methods, including biological control, especially against pentatomid bugs (Eilenberg et al. 2001; Conti et al. 2021; Zapponi et al. 2021). Egg parasitoids are central candidates in biological control strategies, either through conservation biological control of resident parasitoid communities, augmentative releases of native or established species, or classical biological control involving coevolved natural enemies (Eilenberg et al. 2001; Conti et al. 2021; Haye et al. 2015; Zapponi et al. 2021). Among them, Scelionidae, and particularly species of *Trissolcus* Ashmead (1893) and *Telenomus* Haliday (1833), have received considerable attention because several species are closely associated with pentatomid eggs and include key candidates for biological control (Talamas et al. 2015; Talamas et al. 2017; Tortorici et al. 2019; Tortorici et al. 2024).

Other hymenopteran egg parasitoid families are also relevant. In Europe, *Anastatus bifasciatus* (Geoffroy, 1785) (Hymenoptera Eupelmidae) is frequently reported from *H. halys* eggs and has been evaluated for augmentative biological control (Stahl et al. 2019; Iacovone et al. 2022). Finally, species of the genus *Ooencyrtus* Ashmead 1990 (Encyrtidae) has also been reported parasitizing *H. halys* (Fusu & Andreadis, 2023).

However, routine and reliable identification of these parasitoids remains challenging, especially when large numbers of specimens from field surveys must be processed. Recent taxonomic revisions have improved species concepts and identification tools for several groups, particularly within *Trissolcus* and *Telenomus* (Talamas et al. 2015; Talamas et al. 2017; Tortorici et al. 2019; Moraglio et al. 2021a; Tortorici et al. 2024). Morphological identification may nevertheless remain difficult because of incomplete or taxon-specific keys, small body size, sexual dimorphism in some groups (e.g. Eupelmidae), the condition specimens, and, in some taxa, host-induced phenotypic plasticity (Ganjisaffar et al. 2018). As highlighted by several authors (Talamas et al. 2019; Tortorici et al. 2019; Fusu & Andreadis, 2023; Triapitsyn et al. 2020; Potter et al. 2023), molecular approaches including DNA barcoding are therefore useful, and often necessary, to support or confirm morphology-based identifications, process large field samples and detect inconsistencies among morphological determinations, molecular clusters and public reference sequences.

In France, the available information remains scattered among pest-specific surveys and local records, limiting a synthetic view of parasitoid assemblages associated with pentatomid eggs (Bout et al. 2019; Bout et al. 2023).

In this context, the present study aims to clarify the diversity of egg parasitoids associated with four pentatomid species of agronomic interest in France: *Graphosoma italicum* (Müller, 1766)*, H. halys*, *N. viridula*, and *P. prasina*. Additional hosts, including other Pentatomoidea and more distantly related taxa within Panheteroptera *sensu* Wang et al. (2017), were also included to broaden the interpretation of host associations. We combined morphological identification with COI DNA barcoding to assign a species name to specimens, delimit molecular units and identify potential conflicts between morphology, molecular clustering, and public reference sequences. Finally, we discuss the parasitoid species in relation to conservation, augmentative and classical biological control, following the terminology of Eilenberg et al. (2001).

## Materials and Methods

### 1. Sampling

As presented in Table 1, the egg parasitoids characterized in this study came from surveys conducted between 2018 and 2020. While this table summarizes the core sampling campaigns that provided the majority of the specimens, the study also incorporated opportunistic collections across a wider range of settings. Sampling was preferentially performed in agricultural and peri-urban contexts, and the listed habitats and host modalities represent the most frequent contexts encountered. Two methods were used: the collection of naturally-laid egg masses and the exposure of sentinel eggs. For the latter, two host species were used (*H. halys* and *N. viridula*), reared at the National Institute of Agricultural, food and Environment Research (INRAe) Institut Sophia Agrobiotech (ISA), the Interprofessional Technical Center for Fruit and Vegetables (CTIFL) or the National Association of Hazelnut Producers (ANPN), France. Details about the rearing are provided in Bout et al. (2021). *Halyomorpha halys* sentinel egg masses were exposed either directly in the field (hereafter “fresh eggs”), or after storage at cold temperature (-15 °C to -20 °C during at least 48 hours) (hereafter “frozen eggs”). Cold storage is known to kill the embryo within the eggs, facilitating successful development of parasitoids that unable to develop in live eggs (Herlihy et al. 2016; Roversi et al. 2016). Egg masses were glued onto paper strips (1 × 3 cm) and stapled to the underside of leaves of putative host plants of *H. halys* (mostly woody trees, e.g., *Prunus* spp., *Acer* spp.) and *N. viridula* (e.g., horticultural plants). Sentinel egg masses were retrieved within three days. All egg masses were then maintained at Institut Sophia Agrobiotech or ANPN until emergence of parasitoids or nymphs. Each egg mass was placed in a glass tube (1 x 3 cm) with a fine drop of honey and closed with cotton with rearing conditions of 22 ± 1 °C, 60 ± 5% RH, 16:8 h L:D. Tubes were checked daily and emerged parasitoids were preserved in 96% ethanol at −18 °C.

**Table 1:**
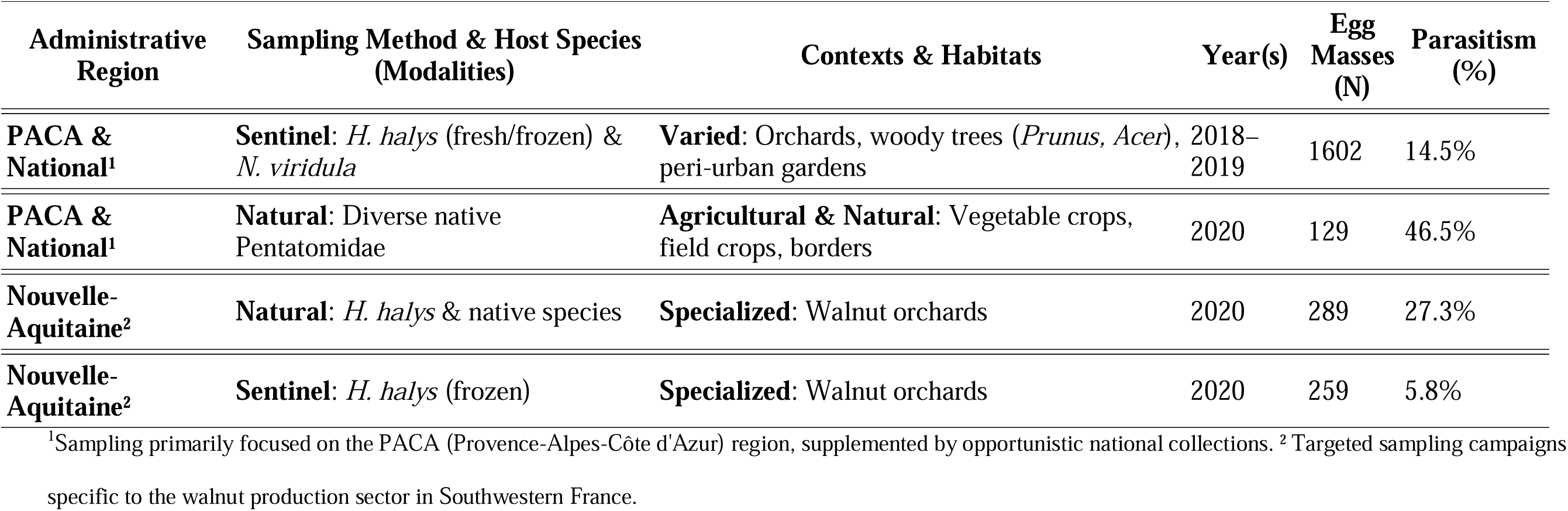
Summary of main sampling effort and parasitism success by region, method, and habitat (2018–2020). Summary of the primary sampling campaigns and parasitism success (adapted from Bout et al. 2021). This table provides a representative overview of the main efforts; it highlights the predominant habitats, crops, and modalities, though additional opportunistic samples and secondary environments were also included in the study.

### 2. Morphological identification

Specimens were identified using keys for *Trissolcus* (Talamas et al. 2015; Talamas et al. 2017; Talamas et al. 2019; Tortorici et al. 2019), *Telenomus* (Tortorici et al. 2024), and *Anastatus* Motchoulsky 1859 (Peng et al. 2020). Ethanol-preserved and DNA-extracted specimens were mounted following Talamas et al. (2017), dried, and card-pointed. Diagnoses were performed with Leica M205C stereomicroscopes (INRAE). Vouchers are deposited at INRAE UMR ISA, Sophia-Antipolis.

## 3. Molecular characterization – DNA Barcoding

### 3.1 DNA extraction and amplification

From each egg mass where parasitoids emerged, three specimens (preferentially females) were selected ad hoc for DNA extraction. Genomic DNA was extracted from ethanol preserved specimens using QuickExtract^TM^ (LUCIGEN MA150E; 30 µL) following the manufacturer’s instructions; this non-destructive protocol preserves vouchers for morphology. A fragment of COI (cytochrome c oxidase subunit I) was amplified with primers LCO1490 and HCO2198 (∼600–700 bp): HCO2198 (50-TAAA CTT CAG GGT GAC CAA AAA ATC A-30), LCO1490(50-GGTC AAC AAA TCA TAA AGA TAT TGG-30) (Folmer et al 1994). Amplicons were Sanger-sequenced by Genewiz (Beckman Coulter Genomics). Residual DNA is archived at INRAE Sophia-Antipolis. Molecular studies were conducted at the National Institute of Agricultural, food and Environment Research (INRAe), Sophia-Antipolis, France.

### 3.2 Haplotype identification

Raw COI sequences were edited and aligned in BioEdit, MEGA X, and Geneious R10. Raw sequences’ ends were truncated to generate fragments of uniform length with unambiguous ends. Haplotypes were defined as unique sequences (100% identity) and enumerated in DnaSP v5.10.01 (Librado & Rozas, 2009). Each haplotype received an internal identifier (Hap_001, Hap_002, …). The sample–haplotype matching is provided in Supplementary Table 1.

To limit NUMTs, pseudogenes and low-reliability sequences, we translated sequences using the invertebrate mitochondrial code and removed sequences with internal stop codons or frameshifting indels. As a stringent precautionary filtering step, singleton COI haplotypes represented by a single individual from a single locality were excluded from downstream clustering and taxonomic interpretations. These haplotypes were retained in Table 3 (Supplemental data) but were not deposited in GenBank. The final haplotype sequences were deposited in GenBank under accession numbers QB009317–QB009404, QB001854–QB001874 and QB001849– QB001853 for Scelionidae, Encyrtidae and Eupelmidae (*Anastatus*) respectively (Table 2).

**Table 2:** Consolidated COI haplotypes and molecular assignments of egg parasitoids collected in France. This table summarizes the molecular diversity across the three studied families (Scelionidae, Eupelmidae, and Encyrtidae). It lists the unique COI haplotypes retained after filtering out singletons and sequences with quality issues, such as internal stop codons. For each Molecular Operational Taxonomic Unit (MOTU), the total number of individuals (N), a representative reference specimen ID, and the corresponding accession numbers for GenBank and/or BOLD are provided.

**Table 3.** Supplemental Data (Annexes): Comprehensive dataset of sampled individuals, collection metadata, and host-parasitoid associations. This supplemental table provides a detailed record for all specimens processed in this study. For each individual, it includes collection dates, geographical locations (administrative regions and/or coordinates), and specific habitat contexts ranging from specialized orchards to peri-urban gardens. It also details the host associations, specifying the host species and egg modalities (natural egg masses versus fresh or frozen sentinel eggs). Finally, it provides the cross-reference between each individual sample and its assigned COI haplotype as listed in Table 2.

Each haplotype was queried against BOLD Systems (ID Engine) and NCBI GenBank (BLASTn) (Altschul et al. 1990). We recorded top hits (percent identity; aligned coverage), ≥ 98–99% identity and ≥ 90% coverage were considered compatible with species-level assignment. The resulting consolidated haplotypes (Table 2) were used for downstream analyses.

### 3.3 Delimitation of molecular operational units (MOTUs)

To delimit Molecular Operational Taxonomic Units (MOTUs), we used an integrative approach combining our dataset with reference sequences from BOLD, GenBank, and literature (Talamas et al. 2019; Tortorici et al. 2019). For each family, sequence clustering was performed using Automatic Partitioning (ASAP) (Puillandre et al. 2021) based on Kimura-2-parameter (K2P) distance matrices (Kimura, 1980). The optimal partition for each family was selected according to the best ASAP score. To visualize these clusters and assess their stability, Neighbor-Joining (NJ) trees (Saitou & Masatochi, 1987) were inferred in MEGA X (Tamura et al. 2013) with 1,000 bootstrap replicates, using the same K2P model for nucleotides and Poisson-corrected distances for amino-acid sequences. Analyses were rooted using specific outgroups: *Hadronotus bicolor* Ashmead, 1894 (GenBank accessions OQ561918.1–OQ561920.1) for Scelionidae, and a species of *Anagyrus* Howard (Malausa et al. 2016) for Encyrtidae.

To ensure robust taxonomic assignments and a global perspective on molecular diversity, our dataset was supplemented with reference sequences retrieved from public databases (NCBI GenBank and BOLD Systems) and from key taxonomic literature (*e.g.*, Talamas et al. 2019; Tortorici et al. 2019). These external references were used as anchors for species identification and to resolve discrepancies between morphological and molecular data.

## Results

### 1. Generalities

Three families of Hymenopteran parasitoids were present in our sampling (by decreasing order of occurrences): Scelionidae, Eupelmidae and Encyrtidae. Their related results are separately discussed below using a common workflow:

i. Information about the observed molecular diversity at the nucleic- and amino-acid levels
ii. Information about additional molecular resources
iii. Results of ASAP partitioning
iv. Discussion about observed “conflicts”, three types being distinguished:

- Type 1: Conflicts between our own molecular and morphological characterizations
- Type 2: Minor (few sequences involved) conflict between the species’ affiliation of Genbank accessions and the obtained clustering
- Type 3: Major conflicts (several sequences from this study or Genbank) questioning the validity of some previously established species.
v. Main patterns about the final alignment and partition, once the conflicts are resolved
vi. Information about the observed diversity in France
vii. For the most abundant species, their observed host ranges (fresh natural *H. halys* sentinel eggs, frozen *H. halys* sentinel eggs, natural eggs of other species)

### 2. Scelionidae (Platygastroidea)

One hundred and twenty-six haplotypes (1,358 individuals) were retrieved from our samples based on aligned sequences of 537 bp (including one indel of 3 bp, see below). The median number of individuals per haplotype is 3, this value however masked a very high variability. Indeed, the 24 most frequent haplotypes (≥ 10 individuals/haplotype) total 1,075 individuals (79%) while, at the opposite, 38 haplotypes were observed only once. These latter were discarded for the rest of the analysis, as explain previously. At the amino-acid level, 29 haplotypes were observed, the 7 most frequent ones (≥ 50 individuals) totalling 1,049 individuals (77%). 112 sequences of Scelionidae — 4 genera: *Gryon* Haliday (1833), *Hadronotus* Förster (1856)*, Telenomus*, and *Trissolcus* — were added from Genbank to our dataset. Fifteen percent of these sequences were significantly shorter than ours for the area investigated. The alignment was modified to accommodate these new sequences, and a second indel (6 bp) was added for *Gryon* sequences.

Based on this final dataset, the ASAP analysis proposed two quite equivalent partitions with the same ASAP score (4.5) and 36–37 different species. The only difference dealt with 2 haplotypes, Hap_027 (2 locations and 8 individuals) and Hap_188 (1 location and 2 individuals), either included in *Trissolcus colemani* (Crawford) in one partition, either grouped together as an unidentified species in the other. Because several individuals of Hap_027 were morphologically identified as *Tr. colemani* by E. Talamas and A. Bout, we used the most parsimonious partition. Pairwise K2P distances revealed a marked barcoding gap (See Figure 1A). The maximum intraspecific divergence (0.06) was indeed substantially lower than the minimum interspecific divergence (0.09).

**Figure 1:**
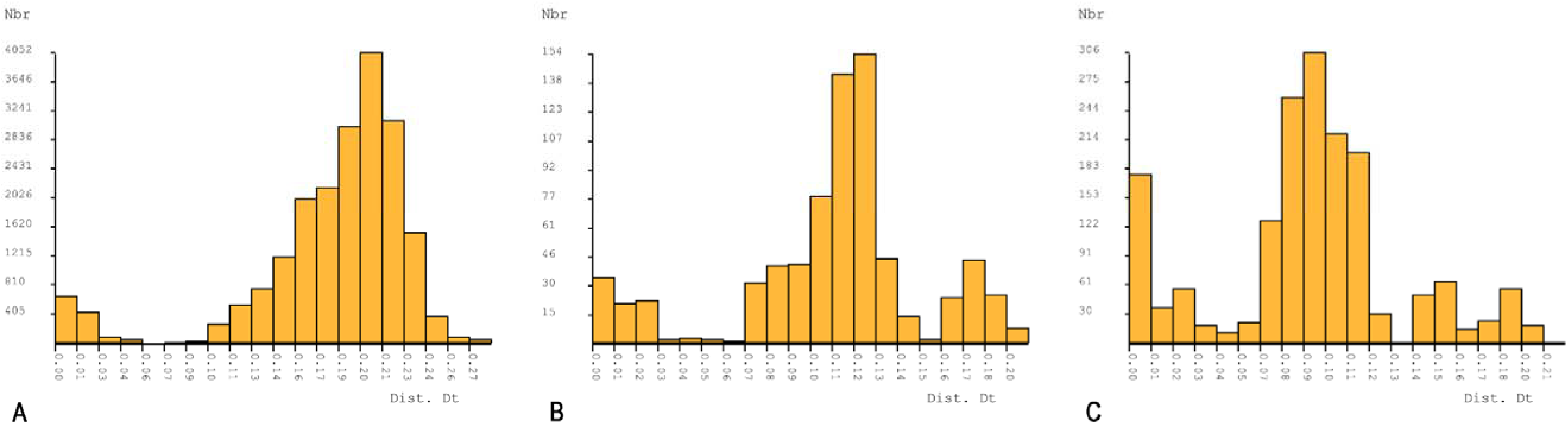
Distribution of pairwise Kimura 2-Parameter (K2P) distances observed during ASAP analysis. Frequency distribution of pairwise COI genetic distances (K2P) for the three studied families: (A) Scelionidae, (B) Eupelmidae, and (C) Encyrtidae. These histograms illustrate the molecular variation used by the Automatic Partitioning (ASAP) algorithm to define species boundaries and MOTUs. The x-axis represents the K2P distance, and the y-axis (Nbr) indicates the number of pairwise sequence comparisons within each family.

**Figure 2:**
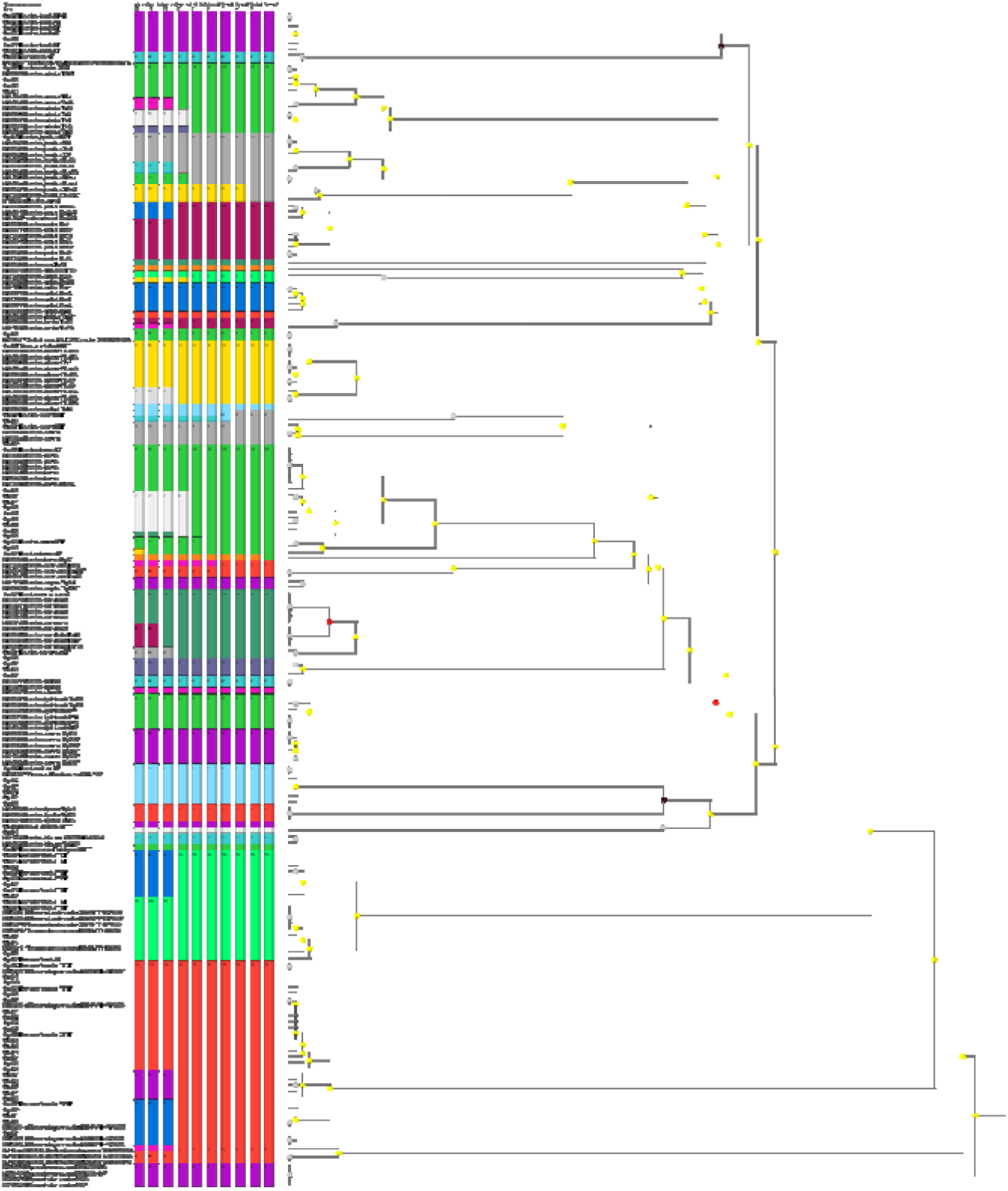
Molecular species delimitation and ASAP partitioning for the family Scelionidae. Detailed results of the Automatic Partitioning (ASAP) analysis for the Scelionidae family, including the genera *Trissolcus* and *Telenomus*. The clusters represent the partition with the optimal ASAP score, defining the Molecular Operational Taxonomic Units (MOTUs) for the study. Specific annotations highlight resolved taxonomic complexes and conflicts, such as for *Tr. viktorovi* complexes, as well as the distinction between the *T. colemani* 1 and 2 lineages. French haplotypes are shown alongside international reference sequences from GenBank and BOLD to support molecular assignments.

Based on this partition, several conflicts were observed. Only one “Type 1” conflict was observed, two haplotypes (Hap_083 and Hap_085) were identified as *Tr. colemani* but clustered with *Trissolcus belenus* (Walker). Two “Type 2” conflicts were observed. Indeed, two Genbank accessions (MN615627 and MN615633) previously identified as *Trissolcus japonicus* (Ashmead) clustered with one sequence of *Trissolcus kozlovi* Rjachovskij (MT345600) and formed a separate cluster in our partition. This pattern may reflect misidentification of older material deposited before the diagnostic separation of *Tr. kozlovi* was further clarified, rather tan uncertainty in the current species concepts(Moraglio et al. 2021b). Moreover, one Genbank accession (MN615643) previously identified as *Tr. belenus* clustered as another unidentified species in our partition. More significantly, 3 “Type 3” conflicts were also observed. First, 6 Genbank accessions (MN615648–MN615653) previously identified as *Tr. colemani* constituted an independent and unidentified cluster. It will be named hereafter *Tr. colemani* 2 as opposed to *Tr. colemani* 1 represented by Hap_027, Hap_061, Hap_188, Hap_189, MK906051 and MN603801. It appears that *Tr. colemani* 1 is related to specimens sampled in Iran (MK906051 and MN603801) (Tortorici et al. 2019) and Europe (this study) while *Tr. colemani 2* is related to individuals sampled in Korea and Japan (Talamas et al. 2019). Second, two sequences, our haplotype Hap_052, initially identified in the working dataset as *Trissolcus simoni* (Mayr), and GenBank accession MN044340, identified as *Trissolcus viktorovi* Kozlov, cluster together. Because *Tr. simoni* is currently considered a junior synonym of *Tr. scutellaris* (Thomson) (Talamas et al. 2017), and because the identification of MN044340 as *Tr. viktorovi* is more recent and based on the revised framework for Paleartic *Trissolcus*, we refer to this cluster as *Tr. viktorovi* hereafter. This cluster was distinct from the *Tr. scutellaris* cluster recovered in our analyses. Given the subtle morphological differences between *Tr. scutellaris* and *Tr. viktorovi* misidentification of older material cannot be excluded. Thirdly, sequences previously identified as *Telenomus heydeni* and *Telenomus truncatus* clustered together in our partition, consistently with the recent revision of Tortorici *et al*. (2024) who treated *Te. heydeni* as a junior synonym of *Te. truncatus*. We therefore refer to this cluster as *Te. truncates* hereafter.

Taken as a whole and contrarily to the 3 other genera (*Gryon*, *Hadronotus* and *Trissolcus*), the genus *Telenomus* appears to be paraphyletic with, on one side, a separate lineage encompassing *Telenomus turesis* Walker and as *Te. truncatus* and, on the other, 2 lineages nested within the cluster of *Trissolcus* sequences, this dichotomy being also supported by an indel of 3 bp. Those two lineages are respectively represented by, on one side, Hap_030 and KR792756 and, on the other, by Hap_129.

Based on (i) our sampling in France (Figure 6A et 6B) (ii) the haplotypes observed more than once (1320 individuals) and (iii) our final partition, two genera were observed, *Telenomus* (Te) and *Trissolcus* (Tr). *Telenomus* was represented by 452 individuals, 43 haplotypes and 4 species/complex including (i) *Te. truncatus* (296 individuals and 27 haplotypes), *Te. turesis* (142 individuals and 14 haplotypes) and two unidentified species, provisionally named “sp6” and “sp7” and respectively represented by Hap_030 (12 individuals) and by Hap_129 (2 individuals). About “sp7” (Hap_129), the morphological examination of some specimens suggests that it does not belong to the *podisi* group. *Trissolcus* was represented by 849 individuals, 40 haplotypes and 9 species: *Tr. basalis* (Wollaston), *Tr. belenus*, *Tr. colemani*_1, *Tr cultratus* (Mayr), *Tr. japonicus*, *Tr. mitsukurii* (Ashmead), *Tr. scutellaris* (Thomson), *Tr. semistriatus* (Nees von Esenbeck), and the *Tr. viktorovi* complex.

Among them, *Tr. basalis* was the far most frequently observed species (449 individuals and 7 haplotypes). The three other main species were *Tr. belenus* (89 individuals and 13 haplotypes), *Tr. cultratus* (88 individuals and 4 haplotypes) and *Tr.viktorovi* (71 individuals and 6 haplotypes). The presence of two exotic species was confirmed, *Tr japonicus* (21 individuals and 1 haplotype) and *Tr mitsukurii* (58 individuals and 1 haplotype) (Bout et al. 2021 and Martel et al. 2024).

Our dataset also includes the hosts on which the species were retrieved (cf. Figure 5). Taken as a whole, Scelionidae were almost always only collected on Pentatomidae. In fact, the only exception is “sp6”, one of the two lineages where there is a conflict between morphological (*Telenomus*-like) and molecular approaches (*Trissolcus* cluster), this species having only been found on Scutellaridae.

As shown in Figure 5 and regarding more precisely the two main *Telenomus* species (by decreasing abundance): *Te. truncatus* was collected on the two sampled sub-families of Pentatomidae, Pentatominae and Podopinae (only one species sampled, *G. italicum* on which it is the co-dominant species). Among Pentatominae, it was collected on all identified genera and species, *i.e*. *Carpocoris* sp. (co-dominant parasitoid), *Dolycoris baccarum* (Linnaeus, 1758) (co-dominant parasitoid), *Eurydema* sp., *N. viridula*, *P. prasina* (dominant parasitoid), *Piezodorus lituratus* (Fabricius, 1794), *Rhaphigaster nebulosa* (Poda, 1761) (co-dominant parasitoid) and *H. halys* (whatever the method of sampling and type of eggs: naturally-laid *versus* sentinel, fresh *versus* frozen).

*Telenomus turesis* was also collected on the two sampled sub-families of Pentatomidae, Pentatominae and Podopinae (*G. italicum* on which it is the co-dominant species). In Pentatominae, it was collected only on four identified taxa, *i.e. Carpocoris* sp. (co-dominant species), *D. baccarum* (co-dominant species), *P. prasina,* and *R. nebulosa*.

Regarding more precisely the four main *Trissolcus* species (by decreasing abundance: *Tr. basalis*, *Tr. belenus*, *Tr. cultratus* and *Tr. viktorovi*), the observed host ranges were as follows (Figure 5):

- *Tr. basalis* — Only hosts from the sub-family Pentatominae were observed: *Carpocoris* sp., *D. baccarum*, *N. viridula* (dominant parasitoid), *P. prasina*, *R. nebulosa,* and *H. halys* (natural eggs and previously frozen sentinel ones).
- *Tr. belenus* — Only hosts from the sub-family Pentatominae were observed: *N. viridula*, *P. prasina*, *R. nebulosa* (co-dominant parasitoid) and *H. halys* (previously frozen sentinel eggs only).
- *Tr. cultratus* — Only hosts from the sub-family Pentatominae were observed: *P. prasina*, *R. nebulosa* and *H. halys* (previously frozen sentinel eggs only).
- *Tr. viktorovi* – Hosts from the two sub-families were observed, Pentatominae (*Eurydema* sp, *N. viridula*, *P. prasina*) and Podopinae (*G. italicum*)

### 3. Eupelmidae (Chalcidoidea)

A total of 11 haplotypes (542 individuals) were retrieved from our samples based on aligned sequences of 546 bp for most of them (2 of them being of only 358 bp). The median number of individuals per haplotype is 1, this value however masking a very high variability. Indeed, the two most frequent haplotypes (≥ 10 individuals/haplotype) total 526 individuals (97%) while, at the opposite, 6 haplotypes were observed only once. These latter were discarded for the rest of the analysis. At the amino-acid level, four haplotypes were observed, the most frequent one (≥ 50 individuals) totalling 505 individuals (93%).

A total of, 33 sequences of Eupelmidae (2 genera: *Anastatus* and *Eupelmus*) were added from Genbank to our dataset, all these sequences had the same length as ours for the area investigated.

Based on this final dataset, the ASAP analysis proposed one best partition with an ASAP score of 3 and 12 different species. Pairwise K2P distances revealed a marked barcoding gap (Figure 1B). The maximum intraspecific divergence (0.06) was substantially lower than the minimum interspecific divergence (0.7), yielding a gap of 0.01. No overlap between intra- and interspecific distance distributions was detected.

Based on this partition, 2 “Type 3” conflicts were observed (Type by Type as defined at the beginning of the Results section and from top to bottom in Figure 3). First, one of the *Anastatus shichengensis* Sheng & Wang, 1997, (Genbank accession OR473693) was indeed separated from the other *A. shichengensis* ones. Few information is however available about all these accessions. Second, the unique sequence of *Anastatus formosanus* Crawford, 1913 (PQ5810) and the two unique sequences of *Anastatus dexingensis* Sheng & Wang, 1997 (PV844027 and PV844032) are clustering into a single cluster (here after dexingensis-formosanus complex). Few information is however available about all these accessions. An additional issue is the case of *Anastatus bifasciatus* (Geoffroy, 1785) only represented by the Genbank accession MW996607 in our partition. This sequence indeed appeared differentiated from some of our haplotypes (Hap 091, Hap 195, Hap 204, Hap 207 and Hap 208) for which we obtain, after this study, certainty of their affiliations to *A. bifasciatus*. It is however beyond the scope of this article to detail this evidence (Bout et al. In prep.).

**Figure 3:**
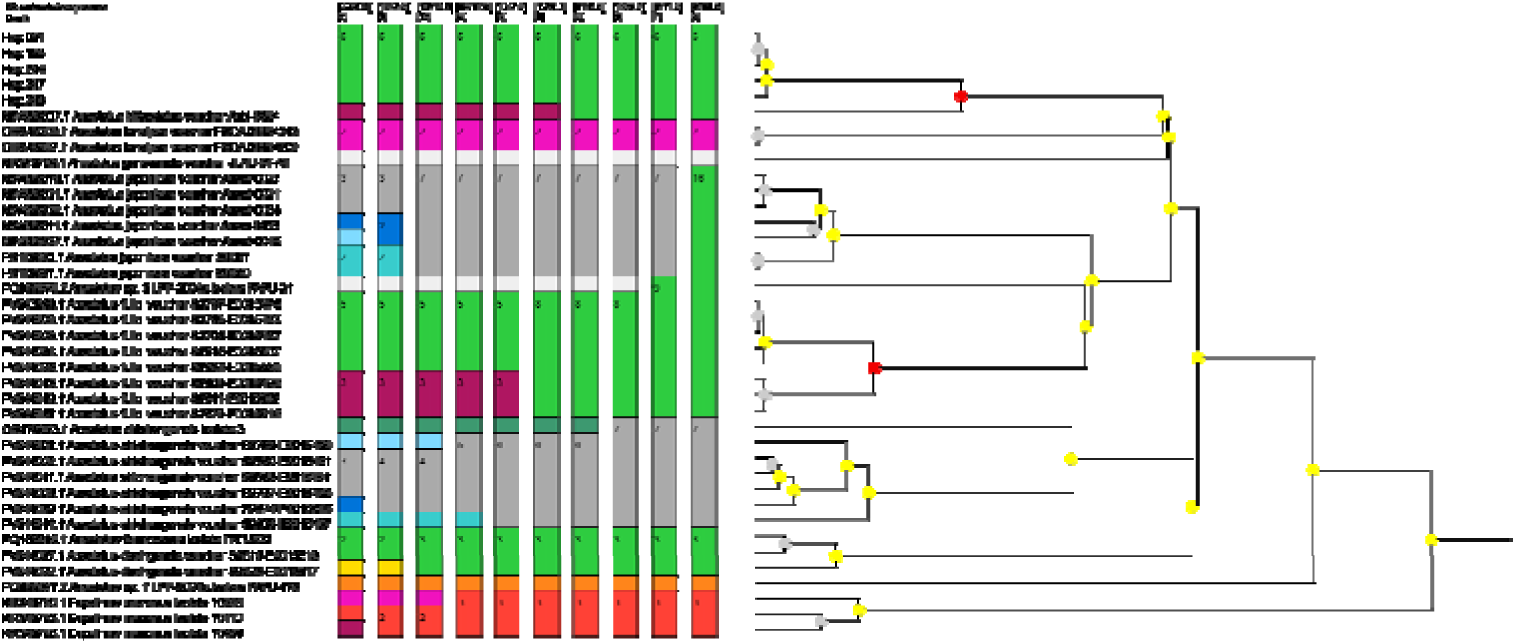
Molecular species delimitation and ASAP partitioning for the family Eupelmidae. Molecular operational taxonomic units (MOTUs) for the Eupelmidae family as defined by the ASAP algorithm. The partitioning primarily focuses on the genus Anastatus, illustrating the molecular proximity between French specimens and public reference sequences. The figure highlights observed Type 3 conflicts, such as the *dexingensis-formosanus* complex, and confirms the status of *Anastatus bifasciatus* as the predominant species in the French sampling. Clusters are ranked according to their ASAP scores to ensure the most robust species delimitation.

Based on (i) our sampling in France (figure 6B), (ii) the haplotypes observed more than once (536 individuals) and (iii) our final partition, only *A. bifasciatus* was observed with 5 different haplotypes.

As shown in Figure 5, *A. bifasciatus* was collected on various hosts of the family Pentatomidae — Pentatominae *(Carpocoris* sp.*, D. baccarum, Eurydema ornata* (Linnaeus, 1758)*, H. halys, N. vidirula, P. prasina*, and *R. nebulosa*), and Podopinae (*G. italicum).* Out of the Pentatomidae, *A. bifascitus* was also collected on Scutelleridae — *Solenosthedium bilunatum* (Lefebvre, 1827), Coreidae — *Gonocerus acuteangulatus* (Goeze, 1778), and Reduvidae.

### 4. Encyrtidae (Super-family: Chalcidoidea)

Twenty-eight haplotypes (343 individuals) were retrieved from our samples based on aligned sequences of 546 bp. The median number of individuals per haplotype is 3, this value however masking a very high variability. Indeed, the seven most frequent haplotypes (≥ 10 individuals/haplotype) total 287 individuals (84%) while, at the opposite, seven haplotypes were observed only once. These latter were discarded for the rest of the analysis. At the amino-acid level, nine haplotypes were observed, the most frequent one (≥ 50 individuals) totalling 270 individuals (79%).

A total of 34 sequences of Encyrtidae (one genus: *Ooencyrtus*) were added from Genbank to our dataset. 79% of these sequences were significantly shorter than ours for the area investigated. Four sequences of *Anagyrus* Howard, 1896 (Encyrtidae) were added as outgroups.

Based on this final dataset, the ASAP analysis proposed one best partition with an ASAP score of 2.5 and 11 different species. Pairwise genetic distances (Figure 1C) showed a multimodal distribution, with a small peak at low divergence values (0 to 0.02) corresponding to intraspecific comparisons and a broader distribution centred around 0.08 to 0.12. However, no clear barcoding gap was detected, as the upper range of intraspecific distances approached the lower range of interspecific distances.

Based on this partition, two “Type 3” conflicts were observed (from top to bottom in Figure 4). Firstly, our partition separated Genbank accessions previously identified as *Ooencyrtus telenomicida* (Vassiliev, 1904) in three distinct clusters. A first one includes specimens from Italy and Israel and match to *O. telenomicida* (Mediterranean populations *sensu* Fusu and Andreadis 2023). The two other clusters observed in our partition distinguish individuals from Romania with, on one side, MN945949 and MN945950 and, on the other, MN945951. This pattern was already observed in Fusu and Andreadis (2023), the authors however considering all these specimens as a single species. It is finally noteworthy that, based on our ASAP analysis, the unique partition grouping all these Romanian specimens is ranked in fourth position with an ASAP score of 4.5.

**Figure 4:**
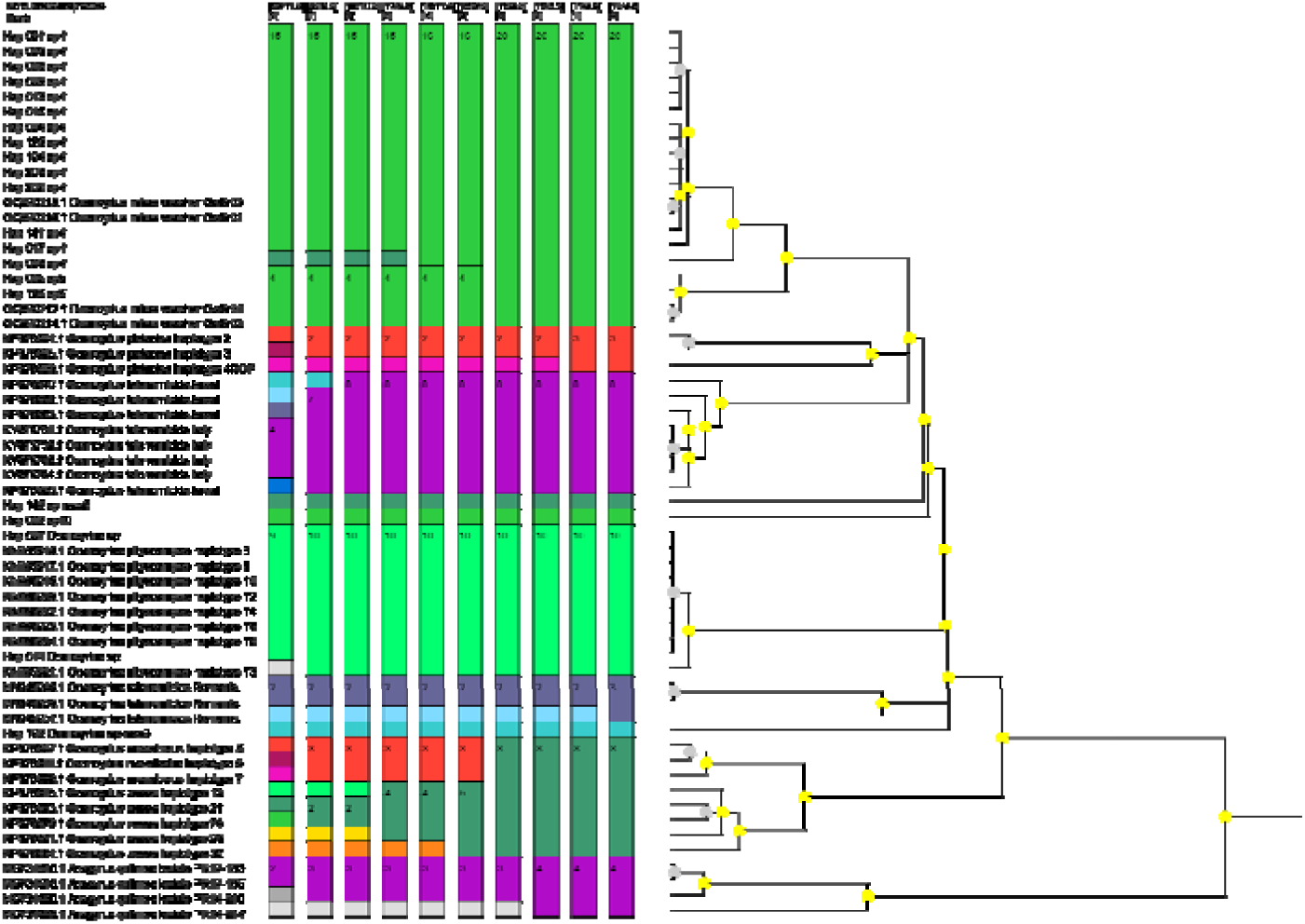
Molecular species delimitation and ASAP partitioning for the family Encyrtidae. Summary of the ASAP partitioning and species boundaries for the Encyrtidae family (genus Ooencyrtus). The figure displays the distribution of unique COI haplotypes into MOTUs, based on the optimal partition score. Key findings include the separation of accessions previously identified as *Ooencyrtus telenomicida* into distinct geographical clusters and the identification of the *mevalbelus-zoeae* complex. This molecular framework supports the morphological characterization of *O. mirus* as the most frequent Encyrtid egg parasitoid retrieved in the survey.

Secondly, all the Genbank accessions affiliated to *Ooencyrtus mevalbelus* Guerrieri & Samra 2018 and *Ooencyrtus zoeae* Guerrieri & Samra, 2018 are clustering in our partition into a single species (hereafter mevalbelus-zoeae complex). These two species seem however actually separated based on multi-locus approach and crossing experiments (Samra et al. 2018 and Fusu and Andreadis 2023). It is finally noteworthy that, based on our ASAP analysis, the unique partition separating these two species is ranked in fifth position with an ASAP score of 8, this partition being also associated with new sub-clustering in *O. mirus* Triapitsyn & Power 2020, *O. pistaciae* Triapitsyn & Power 2019 as well as in *A. quilmes* Triapitsyn, Logarzo & Aguirre 2014.

Based on (i) our sampling in France (figure 6D), (ii) the haplotypes observed more than once (336 individuals) and (iii) our final partition, only one genus, *Ooencyrtus,* was observed with five species: *O. mirus*, *O pityocampae* (Mercet, 1921) and three unidentified species. Among them, *O. mirus* is the most frequently observed species (267 individuals and 16 haplotypes). The two other main species were two of the unidentified species (respectively represented by Hap_142 and Hap_002) with respectively 31 and 23 individuals.

**Figure 5:**
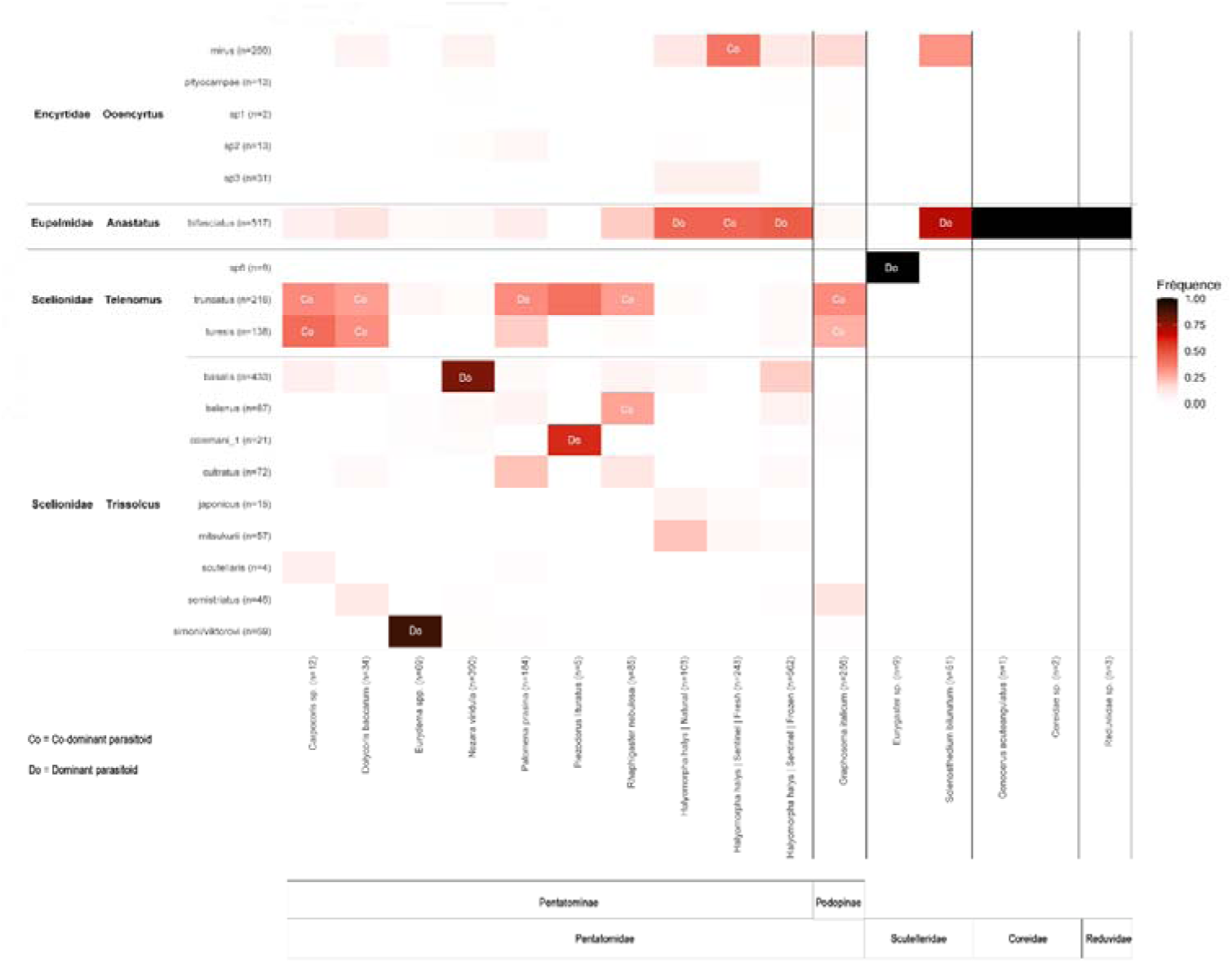
Heatmap of host-parasitoid associations and relative dominance. Matrix summarizing the ecological interactions between the sampled Hemipteran hosts (rows) and the identified egg parasitoid MOTUs (columns). The intensity of the associations is indicated by specific markers: “Do” represents the dominant parasitoid species for a given host, while “Co” indicates co-dominant associations. Molecular Operational Taxonomic Units (MOTUs) correspond to the assignments detailed in Table 2. Empty cells indicate that no parasitism was recorded for that specific host-parasitoid pair during the study.

**Figure 6:**
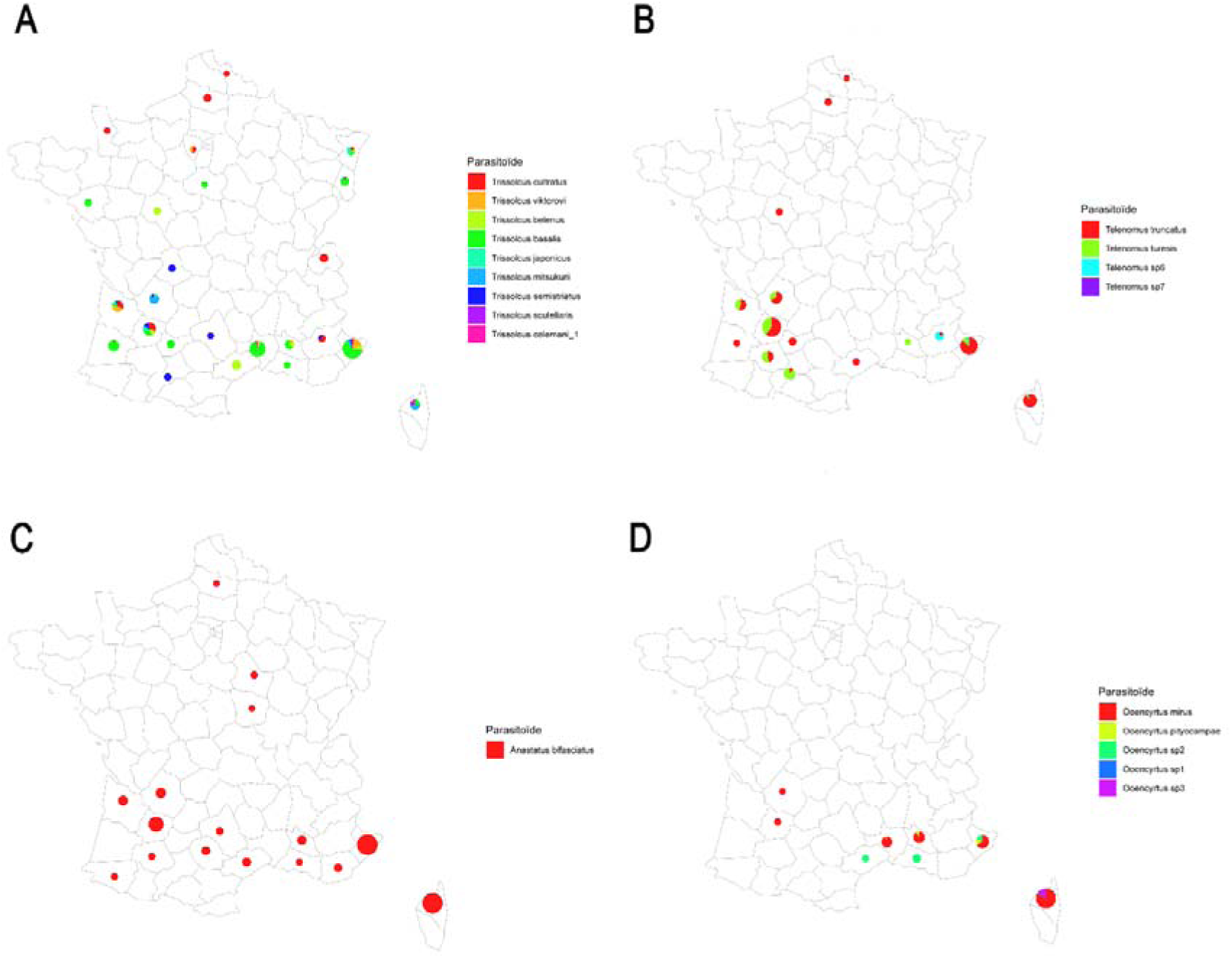
Geographical distribution and species composition of the main egg parasitoid groups in France. Legend: Maps showing the recorded presence and relative abundance of (A) the genus Trissolcus (Scelionidae), (B) the genus Telenomus (Scelionidae), (C) the family Eupelmidae (primarily Anastatus bifasciatus), and (D) the family Encyrtidae (primarily genus Ooencyrtus). Pie charts represent the species or Molecular Operational Taxonomic Units (MOTUs) composition within each French department, as identified in the respective map legends. The diameter of each pie chart is proportional to the total number of individuals successfully sequenced for that group in the department. It should be noted that sampling effort was not standardized across the national territory, with a primary focus on the PACA and Nouvelle-Aquitaine regions.

Regarding *O. mirus*, this species was observed parasitizing hosts of the two sub-families of Pentatomidae, Pentatominae (*D. baccarum*, *N. viridula, H. halys)* and Podopinae (*G. italicum)* as well as one species of Scutelleridae (*S. bilunatum)* (Figure 5).

## Discussion

To our knowledge, the present study represents one of the largest efforts to characterize egg parasitoids of pentatomid bugs using systematic COI DNA barcoding, with approximately 2,000 individuals molecularly assigned across multiple host species in a single country-level survey. While previous monitoring programmes have involved large numbers of egg masses or sampling sites (e.g. Moraglio et al. 2020a; Zapponi et al. 2021; Tortorici et al. 2023), these relied primarily on morphological identification and focused mainly on *Halyomorpha halys*. The combination of taxonomic breadth (multiple host species), sample size for molecular analysis, and integrative morpho-molecular approach distinguishes the present dataset from earlier European surveys.

Obviously, the sampling is clearly not exhaustive since about 80% of the individuals were collected from only four species of the Pentatomidae, *H. halys*, *G. italicum*, *N. viridula* and *P. prasina*. However, several more or less taxonomically related other species were also sampled: five species from the sub family Pentatominae within Pentatomidae, two species from another family (Scutelleridae) within the super-family Pentatomoidea, 3 species from families outside the Pentatomidea (Coreidae and Reduvidae). In the frame of host range assessment, this taxonomic coverage was relevant in a perspective of centrifugal phylogenetic approach (Wapshere 1974).

Species delineation of egg parasitoids was performed here using an integrative approach and, more precisely, by coupling a morphological and molecular characterization. With regard to this latter, a classical DNA barcoding approach was implemented using part of the mitochondrial gene COI as a unique marker (Hebert et al. 2003).

Even if this approach is now popular and routinely used, several limits of this approach have to be kept in mind (Elias et al. 2007, Galtier et al. 2009, Virgilio et al. 2010, Collins & Cruickshank 2012, Cheng et al. 2023). These limits are linked to (i) technical aspects: artefacts generated during sequencing or analysis or incomplete databases, (ii) the nature of the molecular marker: failure to detect interspecific hybridization or (iii) some evolutionary processes: presence of pseudogenes, incomplete lineage sorting, fast speciation with low molecular divergence. This case-study is, of course, no exception to the rule. In particular, two main difficulties were observed. Firstly, some Genbank accessions of Scelionidae were proven invalid after translation with the presence of repeated sequences of amino-acid at the 3’ end or stop codons. Secondly, pseudogenes and technical artefacts are likely to occur within the sequences referred as to *Anastatus bifasciatus* (Bout et al. in prep.). In the present study, singleton haplotypes were not used to support taxonomic conclusions. This choice should not be interpreted as evidence that singleton haplotypes are artefactual, since rare haplotypes may also represent true undersampled diversity.

As often with such studies (Galimberti et al. 2012; Al khatib et al. 2015, Derocles et al. 2016; Tortorici et al. 2019), several potential new taxonomic issues were detected. All were detailed in the paragraph “Results” but we summarize here the most significant ones (Type III). Within Scelionidaethe specimens identified as *Tr. colemani* may represent two cryptic species. The identification of *Tr. simoni* (HQ447084.1) was made prior to the existence of reliable identification tools for Palearctic *Trissolcus* and is likely incorrect. Quite similarly, the same patterns were observed within Eupelmidae with, on one side, a possible sub-division into two species within *A. shichengensis* and, on the other, a possible synonymy between *A. dexingensis* and *A. formosanus*. All these observations would, of course, need further investigations before being validated.

Using an integrative approach, this study provides an up-date of the egg parasitoids associated to the four mains current Pentatomidae pests in France: *H. halys*, *G. italicum*, *N. viridula*, and *P. prasina*. A clear dichotomy appears between, on one side, the three native species (*G. italicum*, *N. viridula* and *P. prasina*) and, on the other, the invasive one (*H. halys*). For the three native species, their guild of egg parasitoids are indeed dominated by one or two Scelionidae species (*Tr*. *basalis* for *N. viridula*, *Te*. *truncatus* for *P. prasina* and the pair of *Te*. *truncatus* and *Te*. *turesis* for *G. italicum*), the same trend being observed for all the additional Pentatomidae species (*Carpocoris* sp., *D. baccarum*, *Eurydema* sp., *P. lituratus* and *R. nebulosa*). Within the Scelionidae, the two main *Telenomus* species (*Te*. *truncatus* and *Te*. *turesis*) appear to be (co-)dominant on several hosts while the native *Trissolcus* (*Tr. basalis*, *Tr. belenus*, *Tr. colemani* 1 and *Tr. viktorovi*) are only (co-) dominant on a single species (respectively, *N. viridula*, *R. nebulosa*, *P. lituratus* and *Eurydema* sp.). Because the sampling efforts are comparable between the *Telenomus* and *Trissolcus* species, these patterns are unlikely to be artefacts and must rather be related to different eco-evolutionary processes linked to ecological specializations and adaptations to the different hosts.

Contrarily to what is observed on native Pentatomidae, the cortege of egg parasitoids observed on the exotic *H. halys* is dominated by the *A. bifasciatus*. This pattern is consistent with several European studies reporting this species as one of the most common, abundant or widespread native egg parasitoids associated with *H. halys* (Moraglio et al. 2020; Moraglio et al. 2021a; Sabbatini Peverieri et al. 2020; Andreadis et al. 2021; Rot et al. 2021; Zapponi et al. 2021). *Anastatus bifasciatus* is of particular interest as it effectively exploits naturally laid egg masses of *H. halys* and was the most frequent species retrieved from this host in our survey.

Cumulated observations about this species paint a picture of a hyper-generalist species with reported parasitism on highly diverse hosts including Hemiptera (in this study, three families within Panheteroptera) (Stahl et al. 2018; Stahl et al. 2019a; Tortorici et al. 2025) but also various Lepidopteran species: *Lymantria dispar* (Linnaeus, 1758) and the pine processionary moth, *Thaumetopoea pityocampa* (Denis & Schiffermüller, 1775) (Masutti, 1964; Stahl et al. 2018; Georgiev et al. 2021) including, at least, a protected one: *Iphiclides podalirius* (Linnaeus, 1758) (Lepidoptera: Papilionidae) in the Var department (Muru, 2021). As *I. podalirius* is a regulated and protected species on the European Red List, these findings highlight the risk of increased contact between the parasitoid and non-target taxa. Such results underscore the need for caution regarding the large-scale spatial and temporal use of *A. bifasciatus* in biocontrol strategies, even when using indigenous strains, and call for rigorous environmental monitoring. As a consequence of this very large host range, the capacity of *A. bifasciatus* to efficiently regulate *H. halys* through conservation biological control may be limited even if destructive “host-feeding” (Konopka et al. 2017) can also impact *H. halys* population alongside parasitism. In complement, *A. bifasciatus* was thus also considered in a context of augmentation biological control (Haye et al. 2015; Stahl et al. 2018, Stahl et al. 2019). Encouraging results were observed in particular in Northern Italy with a significant increase of the parasitism of *H. halys* by *A. bifasciatus* and no documented unintended effect after quite massive releases (Iacovone et al. 2022). Given in particular the risks of unintended effects, the regulatory status about augmentation biocontrol with *A. bifasciatus* still varies among European countries.

Finally, this study confirms the presence in France of two exotic species, *Tr japonicus* and *Tr mitsukurii* that were previously considered as candidates for introduction biological control. In both cases, their introduction is fortuitous, a phenomenon already reported elsewhere, with adventive populations of *Tr. japonicus* detected in North America and Europe, and *Tr. mitsukurii* reported from several European countries (Sabbatini Peverieri et al. 2018; Stahl et al. 2019b; Gariepy et al. 2019; Scaccini et al. 2020; Konjević et al. 2024). At present, the distribution and abundance of these two species are still limited in France but this situation may now evolve rapidly because of the intrinsic dynamics of the already established populations and the deliberate introductions that are now planned. Regarding this last point, the situation in France is now moving from passive surveillance of adventive *Trissolcus* populations towards regulated experimental releases. The use of field-detected French strains of both *Tr. japonicus* and *Tr. mitsukurii* has been authorised for applied trials. First small-scale releases of *Tr. mitsukurii* were conducted in 2024–2025 in Nouvelle-Aquitaine, including hazelnut, kiwifruit and protected eggplant systems whereas moderate releases of *Tr. japonicus* are planned from 2026 onwards, with a stronger regional focus in Nouvelle-Aquitaine. These actions will be accompanied by monitoring of the establishment, parasitism of *H. halys*, interactions with naturally established populations, and possible non-target effects. In protected crops, the approach should be considered mainly as augmentative biological control, whereas outdoor perennial systems may also allow local acclimatization. Future studies should investigate long-term dynamics of the two species from agronomic to ecological perspectives. In addition, the final distribution areas of the two species in France may differ as a consequence of differences in climatic requirements (see for instance, contrasted results obtained in Italy and Turkey: Mele et al. 2026; Ozdemir et al. 2025). The modalities of interactions of the two species in areas of sympatry are also of course of interest with possible outcomes ranging from niche specialization to strong competition for a common resource. Of course, the overall consequences on *H. halys* populations (mean regulation as well as spatio-temporal variability) will be critical to evaluate of biological control. Finally, the possible actual impacts of these species on non-target species will have to be documented, *Tr. mitsukurii* being apparently less-specialized than *Tr. japonicus* (Haye et al. 2020, 2024; Giovannini et al. 2022; Bout et al. unpublished data).

## Supporting information

Table 2 haplotypes

Table 3 supplemental all specimens

## Statements and declarations

The authors have no competing interests to declare.

## Author contributions

Conceptualization: AB, NR; Methodology: AB, RH, NR; Formal analysis and investigation: AB, SW, LC, CC, RH, ET, FT, NR; Writing - original draft preparation: AB, NR; Writing - review and editing: AB, RH, ET, FT, NR; Funding acquisition: AB, RH; Resources: AB, RH; Supervision: AB

## Data availability

The data supporting the findings of this study are provided within the article and its Supplementary Information. High-resolution figures, Supplementary Table S2, and the complete specimen-level dataset are available with the bioRxiv preprint. Representative COI haplotype sequences have been deposited in GenBank under the accession numbers listed in Supplementary Table S2. Additional data are available from the corresponding author upon reasonable request.

## Acknowledgments

We acknowledge all the students, short-term staff, and technical contributors who helped at different stages of this study, including Isabelle LeGoff, Nicolas Bonneti, Lucas Browet, Sarah Diamma, Sophie Dordonnat, Marielena Lahoreau, Alice Leboulanger, Jeremy Desplanques, Guillaume Martel, Amélie Montero, Élodie Rosinsky, Elodie Garnier and Cassandre Vidale, among others. We also thank technical institutes and plant health networks, field partners, growers and occasional contributors who provide samples, field observations or local information including but not limited to Agricultural Chambers, Association Nationale des Producteurs de Noisette (ANPN), Association de Recherche et d’Experimentation sur Fruits et Légumes en Corse (AREFLEC), the Interprofessional Technical Centre for Fruits and Vegetables (CTIFL), Fédération Régionale de Défense contre les Organismes Nuisibles FREDON, and association INVENIO, the experimental orchards of La Pugère and La Morinière, Koppert and Bioplanet. Elijah Talamas was supported by the Florida Department of Agriculture and Consumer Services, Division of Plant Industry.

## References

Alkarrat, H., Kienzle, J., & Zebitz, C. (2020). Biology, abundance and control strategy of Pentatoma rufipes L. (Hemiptera, Pentatomidae) in organic pome fruit orchards in Germany. In: 19th International Conference on Organic Fruit-Growing, Hohenheim/Germany, 17 to February, 19, 2020 2020. pp 111–117

Al Khatib, F., Fusu, L., Cruaud, A., Gibson, G., Borowiec, N., Rasplus, J.-Y., Ris, N., & Delvare, G. (2015). Availability of eleven species names of Eupelmus (Hymenoptera, Eupelmidae) proposed in Al khatib et al. (2014). ZooKeys, 505, 137–145. 10.3897/zookeys.505.9021

Altschul, S. F., Gish, W., Miller, W., Myers, E. W., & Lipman, D. J. (1990). Basic local alignment search tool. Journal of Molecular Biology, 215(3), 403–410. 10.1016/S0022-2836(05)80360-2

Andreadis, S. S., Gogolashvili, N. E., Fifis, G. T., Navrozidis, E. I., & Thomidis, T. (2021). First Report of Native Parasitoids of Halyomorpha halys (Hemiptera: Pentatomidae) in Greece. Insects, 12(11), 984. 10.3390/insects12110984

Ballanger, Y. & Jouffret, P. (1997). La punaise verte et le soja: Un ravageur que l’on peut combattre, même s’ il reste beaucoup à apprendre à son sujet. Phytoma, la défense des végétaux:32–34

Beliën, T., Peusens, G., Schoofs, H., & Bylemans, D. (2015). Stink Bugs (Hemiptera: Pentatomidae) in Pear Orchards: Species Complex, Population Dynamics, Damage Potential and Control Strategies. In: Deckers T, Vercammen J (eds) XII International Pear Symposium, Leuven, Belgium 2015. Acta Hortic, pp 415–420. 10.17660/ActaHortic.2015.1094.53

Blancard, D. (2026). Tomate - Punaises. https://ephytia.inrae.fr/fr/C/5141/Tomate-Punaises.

Bosco, L., Moraglio, S. T., & Tavella, L. (2018). Halyomorpha halys, a serious threat for hazelnut in newly invaded areas. Journal of Pest Science, 91(2), 661–670. 10.1007/s10340-017-0937-x

Bout A., Le Goff I., Cesari L., Genson G., Gard B., Ris N. et Streito J.-C. (2019). Solutions de lutte biologique pour maitriser les punaises. Phytoma 723: 22–27.

Bout, A., Tortorici, F., Hamidi, R., Warot, S., Tavella, L., & Thomas, M. (2021). First Detection of the Adventive Egg Parasitoid of Halyomorpha halys (Stål) (Hemiptera: Pentatomidae) Trissolcus mitsukurii (Ashmead) (Hymenoptera: Scelionidae) in France. Insects, 12(9), 761. 10.3390/insects12090761

Bout A., Iacovone A., Streito JC., Toillon J., Alison B., et Hamidi R. (2023). Solutions et Stratégies pour maîtriser Halyomorpha halys. Phytoma 763: 16–22.

Chen, H.-C. (2023a). A systematic review of the barcoding strategy that contributes to COVID-19 diagnostics at a population level. Frontiers in Molecular Biosciences, 10, 1141534. 10.3389/fmolb.2023.1141534

Chen, H.-C. (2023b). A systematic review of the barcoding strategy that contributes to COVID-19 diagnostics at a population level. Frontiers in Molecular Biosciences, 10, 1141534. 10.3389/fmolb.2023.1141534

Cianferoni, F., Graziani, F., Dioli, P., & Ceccolini, F. (2018). Review of the occurrence of *Halyomorpha halys* (Hemiptera: Heteroptera: Pentatomidae) in Italy, with an update of its European and World distribution. Biologia 73: 599–607. 10.2478/s11756-018-0067-9

Collins, R. A., & Cruickshank, R. H. (2013). The seven deadly sins of DNA barcoding. Molecular Ecology Resources, 13(6), 969–975. 10.1111/1755-0998.12046

Conti, E., Avila, G., Barratt, B., Cingolani, F., Colazza, S., Guarino, S., Hoelmer, K., Laumann, R. A., Maistrello, L., Martel, G., Peri, E., Rodriguez Saona, C., Rondoni, G., Rostás, M., Roversi, P. F., Sforza, R. F. H., Tavella, L., & Wajnberg, E. (2021). Biological control of invasive stink bugs: Review of global state and future prospects. Entomologia Experimentalis et Applicata, 169(1), 28–51. 10.1111/eea.12967

Derocles, S. a. P., Plantegenest, M., Rasplus, J.-Y., Marie, A., Evans, D. M., Lunt, D. H., & Le Ralec, A. (2016). Are generalist Aphidiinae (Hym. Braconidae) mostly cryptic species complexes? Systematic Entomology, 41(2), 379–391. 10.1111/syen.12160

Driss, L., Hamidi, R., Andalo, C., & Magro, A. (2024). Study of the overwintering ecology of the hazelnut pest, *Palomena prasina* (L.) (Hemiptera: Pentatomidae) in a perspective of Integrated Pest Management. Journal of Applied Entomology, 148(1), 34–48. 10.1111/jen.13206

Eilenberg, J., Hajek, A., & Lomer, C. (2001). Suggestions for unifying the terminology in biological control. (46), 387–400.

Elias, M., Hill, R. I., Willmott, K. R., Dasmahapatra, K. K., Brower, A. V. Z., Mallet, J., & Jiggins, C. D. (2007). Limited performance of DNA barcoding in a diverse community of tropical butterflies. Proceedings of the Royal Society B: Biological Sciences, 274(1627), 2881–2889. 10.1098/rspb.2007.1035

Folmer, O., Black, M., Hoeh, W., Lutz, R., & Vrijenhoek, R. (1994). DNA primers for amplification of mitochondrial cytochrome c oxidase subunit I from diverse metazoan invertebrates. Molecular Marine Biology and Biotechnology, 3(5), 294–299.

Fusu, L., & Andreadis, S. S. (2023). Ooencyrtus mirus (Hymenoptera, Encyrtidae), discovered in Europe parasitizing eggs of Halyomorpha halys (Hemiptera, Pentatomidae). Journal of Hymenoptera Research, 96, 1045–1060. 10.3897/jhr.96.109739

Galimberti, A., Romano, D. F., Genchi, M., Paoloni, D., Vercillo, F., Bizzarri, L., Sassera, D., Bandi, C., Genchi, C., Ragni, B., & Casiraghi, M. (2012). Integrative taxonomy at work: DNA barcoding of taeniids harboured by wild and domestic cats. Molecular Ecology Resources, 12(3), 403–413. 10.1111/j.1755-0998.2011.03110.x

Galtier, N., Nabholz, B., Glémin, S., & Hurst, G. D. D. (2009). Mitochondrial DNA as a marker of molecular diversity: A reappraisal. Molecular Ecology, 18(22), 4541–4550. 10.1111/j.1365-294X.2009.04380.x

Ganjisaffar, F., Talamas, E. J., Bon, M.-C., Brown, B. V., Gonzalez, L., & Perring, T. M. (2018). Trissolcus hyalinipennis Rajmohana & Narendran (Hymenoptera, Scelionidae), a parasitoid of Bagrada hilaris (Burmeister) (Hemiptera, Pentatomidae), emerges in North America. Journal of Hymenoptera Research, 65, 111–130. 10.3897/jhr.65.25620

Gard, B., Bout, A., & Pierre, P. (2022). Release strategies of Trissolcus basalis (Scelionidae) in protected crops against Nezara viridula (Pentatomidae): Less is more. Crop Protection, 161, 106069. 10.1016/j.cropro.2022.106069

Gariepy, T. D., & Talamas, E. J. (2019). Discovery of Trissolcus japonicus (Hymenoptera: Scelionidae) in Ontario, Canada. The Canadian Entomologist, 151(6), 824–826. 10.4039/tce.2019.58

Georgiev, G., Rousselet, J., Laparie, M., Robinet, C., Georgieva, M., Zaemdzhikova, G., Roques, A., Bernard, A., Poitou, L., Buradino, M., Kerdelhue, C., Rossi, J.-P., Matova, M., Boyadzhiev, P., & Mirchev, P. (2021). Comparative studies of egg parasitoids of the pine processionary moth (*Thaumetopoea pityocampa*, Den. & Schiff.) in historic and expansion areas in France and Bulgaria. Forestry: An International Journal of Forest Research, 94(2), 324–331. 10.1093/forestry/cpaa022

Giovannini, L., Sabbatini-Peverieri, G., Marianelli, L., Rondoni, G., Conti, E., & Roversi, P. F. (2022). Physiological host range of Trissolcus mitsukurii, a candidate biological control agent of Halyomorpha halys in Europe. Journal of Pest Science, 95(2), 605–618. 10.1007/s10340-021-01415-x

Gomes, E., Houdiard, E., Toillon, J., Thomas, M., & Hamidi, R. (2025). Evaluation of an Attract-and-Kill strategy against stink bugs in French hazelnut orchards. Paper presented at the Conference: XI International Congress on Hazelnut, 4-8 August 2025, Beijing, China,

Hamidi, R., Calvy, M., Valentie, E., Driss, L., Guignet, J., Thomas, M., & Tavella, L. (2022). Symptoms resulting from the feeding of true bugs on growing hazelnuts. Entomologia Experimentalis et Applicata, 170(6), 477–487. 10.1111/eea.13165

Hamidi, R., Rouzes, R., Toillon, J., Thomas, M., & Tavella, L. (2022). The green shield bug, Palomena prasina, and the red-legged shield bug, Pentatoma rufipes, two secondary pests of French hazelnuts? Acta Hortic:1–9

Haye, T., Fischer, S., Zhang, J., & Gariepy, T. (2015). Can native egg parasitoids adopt the invasive brown marmorated stink bug, Halyomorpha halys (Heteroptera: Pentatomidae), in Europe? Journal of Pest Science, 88(4), 693–705. 10.1007/s10340-015-0671-1

Haye, T., Hoelmer, K., Rossi, J.-P., & Streito, J.-C. (2014). Analyse de risque phytosanitaire express Halyomorpha halys—La punaise diabolique. Anses Rapport d’expertise Collective, (FEBRUARY), 90.

Haye, T., Moraglio, S. T., Stahl, J., Visentin, S., Gregorio, T., & Tavella, L. (2020). Fundamental host range of Trissolcus japonicus in Europe. Journal of Pest Science, 93(1), 171–182. 10.1007/s10340-019-01127-3

Haye, T., Moraglio, S. T., Tortorici, F., Marazzi, C., Gariepy, T. D., & Tavella, L. (2024). Does the fundamental host range of Trissolcus japonicus match its realized host range in Europe? Journal of Pest Science, 97(1), 299–321. 10.1007/s10340-023-01638-0

Hebert, P. D. N., Cywinska, A., Ball, S. L., & deWaard, J. R. (2003). Biological identifications through DNA barcodes. *Proceedings*. Biological Sciences, 270(1512), 313–321. 10.1098/rspb.2002.2218

Herlihy, M. V., Talamas, E. J., & Weber, D. C. (2016). Attack and Success of Native and Exotic Parasitoids on Eggs of Halyomorpha halys in Three Maryland Habitats. PLOS ONE, 11(3), e0150275. 10.1371/journal.pone.0150275

Iacovone, A., Masetti, A., Mosti, M., Conti, E., & Burgio, G. (2022). Augmentative biological control of Halyomorpha halys using the native European parasitoid Anastatus bifasciatus: Efficacy and ecological impact. Biological Control, 172, 104973. 10.1016/j.biocontrol.2022.104973

Kimura, M. (1980). A simple method for estimating evolutionary rates of base substitutions through comparative studies of nucleotide sequences. Journal of Molecular Evolution, 16(2), 111–120. 10.1007/BF01731581

Kiritani, K. (2006). Predicting impacts of global warming on population dynamics and distribution of arthropods in Japan. Population Ecology 48:5–12. 10.1007/s10144-005-0225-0

Kiritani, K. (2011). Impacts of global warming on Nezara viridula and its native congeneric species. Journal of Asia-Pacific Entomology 14:221–226

Konjević, A., Tavella, L., & Tortorici, F. (2024). The First Records of Trissolcus japonicus (Ashmead) and Trissolcus mitsukurii (Ashmead) (Hymenoptera, Scelionidae), Alien Egg Parasitoids of Halyomorpha halys (Stål) (Hemiptera, Pentatomidae) in Serbia. Biology, 13(5), 316. 10.3390/biology13050316

Konopka, J. K., Haye, T., Gariepy, T., Mason, P., Gillespie, D., & McNeil, J. N. (2017). An exotic parasitoid provides an invasional lifeline for native parasitoids. Ecology and Evolution, 7(1), 277–284. 10.1002/ece3.2577

Leskey, T. C., & Nielsen, A. L. (2018). Impact of the Invasive Brown Marmorated Stink Bug in North America and Europe: History, Biology, Ecology, and Management. Annual Review of Entomology, 63(1), 599–618. 10.1146/annurev-ento-020117-043226

Librado, P., & Rozas, J. (2009). DnaSP v5: A software for comprehensive analysis of DNA polymorphism data. Bioinformatics, 25(11), 1451–1452. 10.1093/bioinformatics/btp187

Malausa, T., Delaunay, M., Fleisch, A., Groussier-Bout, G., Warot, S., Crochard, D., Guerrieri, E., Delvare, G., Pellizzari, G., Kaydan, M. B., Al-Khateeb, N., Germain, J.-F., Brancaccio, L., Le Goff, I., Bessac, M., Ris, N., & Kreiter, P. (2016). Investigating Biological Control Agents for Controlling Invasive Populations of the Mealybug Pseudococcus comstocki in France. PLOS ONE, 11(6), e0157965. 10.1371/journal.pone.0157965

Martel, G., Bout, A., Tortorici, F., Hamidi, R., Tavella, L., & Thomas, M. (2024). First detection of Trissolcus japonicus (Ashmead) (Hymenoptera, Scelionidae) in southwestern France. Journal of Hymenoptera Research, 97, 1123–1139. 10.3897/jhr.97.132433

Masutti, L. (1964). Ricerche sui parassiti oofagi della Thaumetopoea pityocampa Schiff. Annali Centro Econ Mont Venezie 1963-1964; 4: 205-271, 4, 205–271.

McPherson, J.E. (2018). Invasive Stink Bugs and Related Species (Pentatomoidea): biology, Higher Systematics, Semiochemistry, and Management. CRC Press, Taylor & Francis Group, Boca Raton, USA

Mele, A., Mills, N. J., Canella, J., Mirandola, E., Ceccato, E., Tirello, P., Scaccini, D., Abram, P. K., & Pozzebon, A. (2026). Population-level impact of egg parasitism on Halyomorpha halys despite a rapid shift in parasitoid species composition. Journal of Pest Science, 99(1), 18. 10.1007/s10340-025-01973-4

Moraglio, S. T., Tortorici, F., Pansa, M. G., Castelli, G., Pontini, M., Scovero, S., Visentin, S., & Tavella, L. (2020). A 3-year survey on parasitism of Halyomorpha halys by egg parasitoids in northern Italy. Journal of Pest Science, 93(1), 183–194. 10.1007/s10340-019-01136-2

Moraglio, S. T., Tortorici, F., Giromini, D., Pansa, M. G., Visentin, S., & Tavella, L. (2021a). Field collection of egg parasitoids of Pentatomidae and Scutelleridae in Northwest Italy and their efficacy in parasitizing *Halyomorpha halys* under laboratory conditions. Entomologia Experimentalis et Applicata, 169(1), 52–63. 10.1111/eea.12966

Moraglio, S.T., Tortorici, F., Visentin, S., Pansa, M.G., & Tavella, L. (2021b). *Trissolcus kozlovi* in North Italy: host specificity and augmentative releases against *Halyomorpha halys* in hazelnut orchards. Insects 12: 464. 10.3390/insects12050464

Muru, D. (2021). Magic bullet or shot in the dark? Potential and limits of biological control for experimental ecology. Nice Université Côte d’Azur.

Ozdemir, I. O., Şılbır, M. F., Karadağ, E., Alaybay, A., Özer, G., Tortorici, F., Walton, V. M., & Dogan, F. (2025). Low Natural Parasitism of the Invasive Halyomorpha halys Versus Strong Native Suppression of Palomena prasina: Evidence from a Three-Year Survey in Northwestern Türkiye. Insects, 16(12), 1212. 10.3390/insects16121212

Peng, L., Gibson, G. A. P., Tang, L., & Xiang, J. (2020). Review of the species of Anastatus (Hymenoptera: Eupelmidae) known from China, with description of two new species with brachypterous females. Zootaxa, 4767(3). 10.11646/zootaxa.4767.3.1

Peverieri, G. S., Talamas, E., Bon, M. C., Marianelli, L., Bernardinelli, I., Malossini, G., Benvenuto, L., Roversi, P. F., & Hoelmer, K. (2018). Two asian egg parasitoids of halyomorpha halys (stål) (hemiptera, pentatomidae) emerge in northern italy: Trissolcus mitsukurii (ashmead) and trissolcus japonicus (ashmead) (hymenoptera, scelionidae). Journal of Hymenoptera Research, (67), 37–53. 10.3897/jhr.67.30883

Potter, M., Bremer, J., Moore, M., Talamas, E., & Shrewsbury, P. (2023). Telenomus cristatus Johnson (Hymenoptera, Scelionidae): New diagnostic data, distribution records and host associations. Biodiversity Data Journal, 11, e111347. 10.3897/BDJ.11.e111347

Powell, G. (2020). The biology and control of an emerging shield bug pest, *Pentatoma rufipes* (L.) (Hemiptera: Pentatomidae). Agricultural and Forest Entomology 22:298–308. 10.1111/afe.12408

Puillandre, N., Brouillet, S., & Achaz, G. (2021). ASAP: Assemble species by automatic partitioning. Molecular Ecology Resources, 21(2), 609–620. 10.1111/1755-0998.13281

Rot, M., Maistrello, L., Costi, E., Bernardinelli, I., Malossini, G., Benvenuto, L., & Trdan, S. (2021). Native and Non-Native Egg Parasitoids Associated with Brown Marmorated Stink Bug (Halyomorpha halys [Stål, 1855]; Hemiptera: Pentatomidae) in Western Slovenia. Insects, 12(6), 505. 10.3390/insects12060505

Roversi, Pio Federico. (2016). Searching for native egg parasitoids of the insive alien species halyomorphahalys Stal (heteróptera Pentatomidae) in southern Europe. Redia, 63–70. 10.19263/Redia-99.16.01

Sabbatini-Peverieri, G., Giovannini, L., Benvenuti, C., Madonni, L., Hoelmer, K., & Roversi, P. F. (2020). Characteristics of the meconia of European egg parasitoids of Halyomorpha halys. Journal of Hymenoptera Research, 77, 187–201. 10.3897/jhr.77.52904

Saitou, N., & Masatochi, N. (1987). The neighbor-joining method: A new method for reconstructing phylogenetic trees. Molecular Biology and Evolution, 4(4), 406–425. 10.1093/oxfordjournals.molbev.a040454

Samra, S., Cascone, P., Noyes, J., Ghanim, M., Protasov, A., Guerrieri, E., & Mendel, Z. (2018). Diversity of Ooencyrtus spp. (Hymenoptera: Encyrtidae) parasitizing the eggs of Stenozygum coloratum (Klug) (Hemiptera: Pentatomidae) with description of two new species. PLOS ONE, 13(11), e0205245. 10.1371/journal.pone.0205245

Scaccini, D., Falagiarda, M., Tortorici, F., Martinez-Sañudo, I., Tirello, P., Reyes-Domínguez, Y., Gallmetzer, A., Tavella, L., Zandigiacomo, P., Duso, C., & Pozzebon, A. (2020). An insight into the role of trissolcus mitsukurii as biological control agent of halyomorpha halys in Northeastern Italy. Insects, 11(5), 1–16. 10.3390/insects11050306

Stahl, J. M., Babendreier, D., & Haye, T. (2018). Using the egg parasitoid Anastatus bifasciatus against the invasive brown marmorated stink bug in Europe: Can non-target effects be ruled out? Journal of Pest Science, 91(3), 1005–1017. 10.1007/s10340-018-0969-x

Stahl, J. M., Babendreier, D., Marazzi, C., Caruso, S., Costi, E., Maistrello, L., & Haye, T. (2019). Can Anastatus bifasciatus Be Used for Augmentative Biological Control of the Brown Marmorated Stink Bug in Fruit Orchards? Insects, 10(4), 108. 10.3390/insects10040108

Stahl, J., Tortorici, F., Pontini, M., Bon, M.-C., Hoelmer, K., Marazzi, C., Tavella, L., & Haye, T. (2019). First discovery of adventive populations of Trissolcus japonicus in Europe. Journal of Pest Science, 92(2), 371–379. 10.1007/s10340-018-1061-2

Talamas, E. J., Bon, M.-C., Hoelmer, K. A., & Buffington, M. L. (2019). Molecular phylogeny of Trissolcus wasps (Hymenoptera, Scelionidae) associated with Halyomorpha halys (Hemiptera, Pentatomidae). Journal of Hymenoptera Research, 73, 201–217. 10.3897/jhr.73.39563

Talamas, E. J., Buffington, M. L., & Hoelmer, K. (2017). Revision of Palearctic Trissolcus Ashmead (Hymenoptera, Scelionidae). Journal of Hymenoptera Research, 56, 79–261. 10.3897/jhr.56.10158

Talamas, E. J., Johnson, N. F., & Buffington, M. (2015). Key to Nearctic species of Trissolcus Ashmead (Hymenoptera, Scelionidae), natural enemies of native and invasive stink bugs (Hemiptera, Pentatomidae). Journal of Hymenoptera Research, 43, 45–110. 10.3897/JHR.43.8560

Tortorici, F., Orrù, B., Timokhov, A. V., Bout, A., Bon, M.-C., Tavella, L., & Talamas, E. J. (2024). Telenomus Haliday (Hymenoptera, Scelionidae) parasitizing Pentatomidae (Hemiptera) in the Palearctic region. Journal of Hymenoptera Research, 97, 591–620. 10.3897/jhr.97.127112

Tortorici, F., Talamas, E. J., Moraglio, S. T., Pansa, M. G., Asadi-Farfar, M., Tavella, L., & Caleca, V. (2019). A morphological, biological and molecular approach reveals four cryptic species of Trissolcus Ashmead (Hymenoptera, Scelionidae), egg parasitoids of Pentatomidae (Hemiptera). Journal of Hymenoptera Research, 73, 153–200. 10.3897/jhr.73.39052

Tortorici, F., Bombi, P., Loru, L., Mele, A., Moraglio, S. T., Scaccini, D., Pozzebon, A., Pantaleoni, R. A., & Tavella, L. (2023). Halyomorpha halys and its egg parasitoids *Trissolcus japonicus* and *T. mitsukurii*: The geographic dimension of the interaction. NeoBiota, 85, 197–221. 10.3897/neobiota.85.102501

Tortorici, S., Cavallaro, C., Siscaro, G., Lisi, F., Gugliuzzo, A., Roversi, P. F., Tortorici, F., & Rizzo, R. (2025). The Box Bug Gonocerus acuteangulatus (Hemiptera: Coreidae) and Its Egg Parasitoids: Updates on Biocontrol in a Hazelnut Producing Area in Southern Italy. Insects, 16(12), 1281. 10.3390/insects16121281

Triapitsyn, S. V., Andreason, S. A., Power, N., Ganjisaffar, F., Fusu, L., Dominguez, C., & Perring, T. M. (2020). Two new species of Ooencyrtus (Hymenoptera, Encyrtidae), egg parasitoids of the bagrada bug Bagrada hilaris (Hemiptera, Pentatomidae), with taxonomic notes on Ooencyrtus telenomicida. Journal of Hymenoptera Research, 76, 57–98. 10.3897/jhr.76.48004

Virgilio, M., Backeljau, T., Nevado, B., & De Meyer, M. (2010). Comparative performances of DNA barcoding across insect orders. BMC Bioinformatics, 11(1), 206. 10.1186/1471-2105-11-206

Wang, Y., Wu, H., Rédei, D., Xie, Q., Chen, Y., Chen, P., Dong, Z., Dang, K., Damgaard, J., Štys, P., Wu, Y., Luo, J., Sun, X., Hartung, V., Kuechler, S. M., Liu, Y., Liu, H., & Bu, W. (2019). When did the ancestor of true bugs become stinky? Disentangling the phylogenomics of Hemiptera–Heteroptera. Cladistics, 35(1), 42–66. 10.1111/cla.12232

Wapshere, A. J. (1974). A strategy for evaluating the safety of organisms for biological weed control. Annals of Applied Biology, 77(2), 201–211. 10.1111/j.1744-7348.1974.tb06886.x

Zapponi, L., Bon, M. C., Fouani, J. M., Anfora, G., Schmidt, S., & Falagiarda, M. (2020). Assemblage of the Egg Parasitoids of the Invasive Stink Bug Halyomorpha halys: Insights on Plant Host Associations. Insects, 11(9), 588. 10.3390/insects11090588

Zapponi, L., Tortorici, F., Anfora, G., Bardella, S., Bariselli, M., Benvenuto, L., Bernardinelli, I., Butturini, A., Caruso, S., Colla, R., Costi, E., Culatti, P., Di Bella, E., Falagiarda, M., Giovannini, L., Haye, T., Maistrello, L., Malossini, G., Marazzi, C., … Sabbatini-Peverieri, G. (2021). Assessing the Distribution of Exotic Egg Parasitoids of Halyomorpha halys in Europe with a Large-Scale Monitoring Program. Insects, 12(4), 316. 10.3390/insects12040316

