## Supplementary figures and images for "Diversity of egg parasitoids of stink bugs in France, with emphasis on Scelionidae parasitizing main Pentatomidae pests"

### Fig1_barcoding_gap.tif

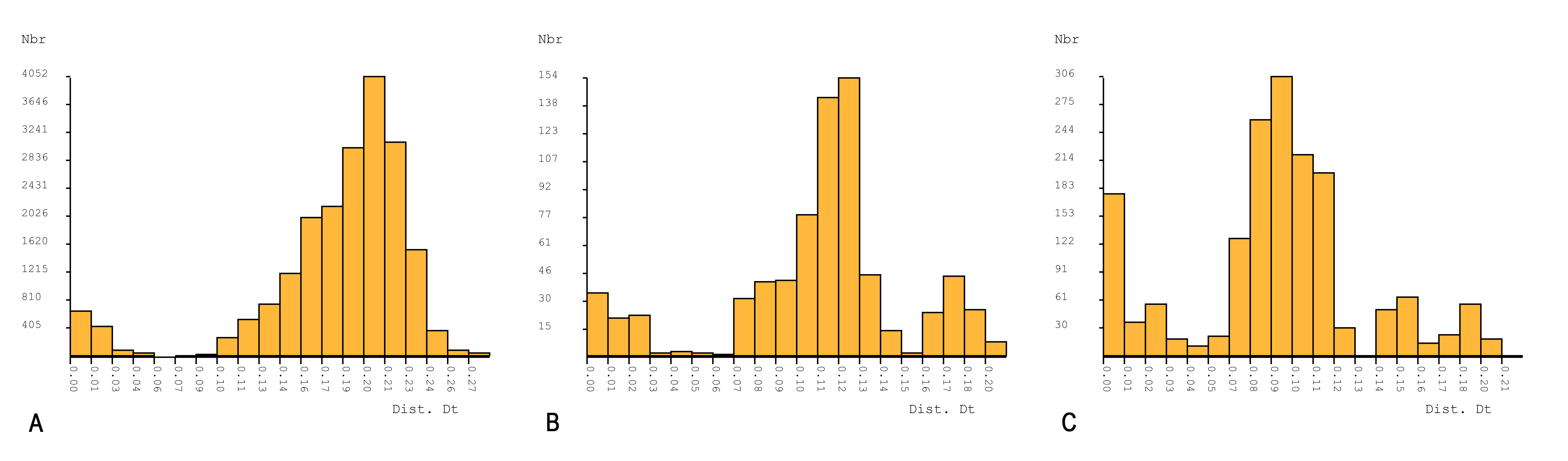

### Fig5_heatmap_host_association.tif

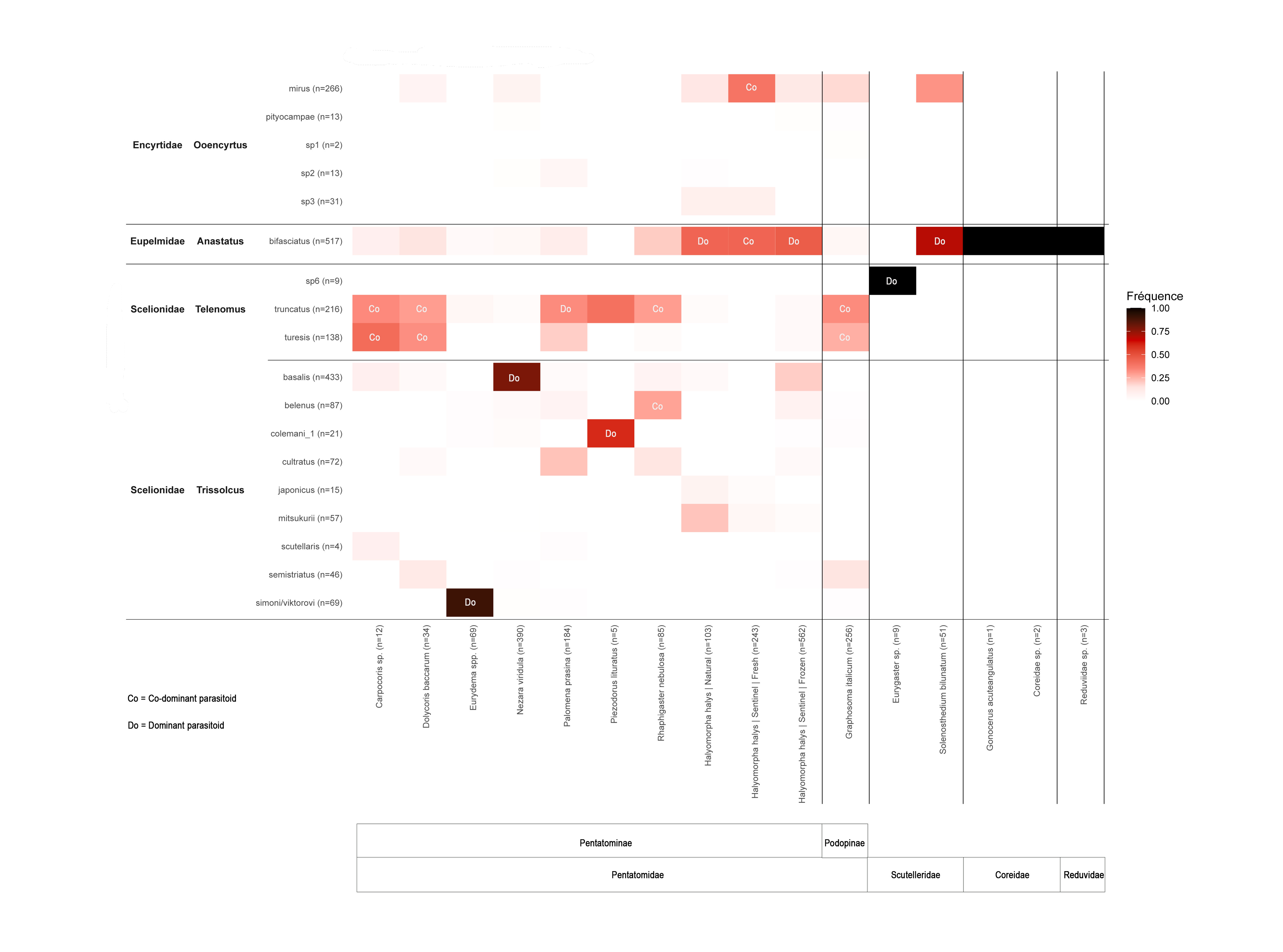
